# *In-vivo* detection of cyclic-di-AMP in *Staphylococcus aureus*

**DOI:** 10.1101/2022.05.16.492224

**Authors:** Nagaraja Mukkayyan, Raymond Poon, Philipp N. Sander, Li-Yin Lai, Zahra Zubair-Nizami, Ming C. Hammond, Som S. Chatterjee

**Author notes:** Equal contributors (Nagaraja Mukkayyan, Raymond Poon). Correspondence: Ming C. Hammond and Som S. Chatterjee.

## Abstract

Cyclic-di-AMP (CDA) is a signaling molecule that controls various cellular functions including antibiotic tolerance and osmoregulation in *Staphylococcus aureus*. In this study, we developed a novel biosensor (*bsuO* P6-4) for *in-vivo* detection of CDA in *S. aureus*. Our study showed that *bsuO* P6-4 could detect a wide concentration range of CDA in both laboratory and clinical strains making it suitable for use in both basic and clinical research applications.

## Main text

Cyclic-di-AMP (CDA) is a newly discovered second messenger, which is present in bacteria belonging to the phyla firmicutes and actinobacteria (1). Recent studies have demonstrated that CDA plays important roles in regulating vital biological processes such as DNA repair, ion homeostasis, and central carbon metabolism among others (2-5). In addition, CDA has also been implicated in controlling processes that are important for bacterial pathogenesis such as biofilm formation, antibiotic tolerance, and virulence (6–10). CDA mediates its function through binding to its cognate effectors (i.e. proteins and riboswitches) and thereby modifying their function through allosteric changes and/or through altered gene expression. Thus, maintaining the correct concentration of CDA in bacterial cells is critical not only to retaining cellular homeostasis but also to responding to changing environmental needs. This is attained by controlling CDA’s synthesis or degradation machinery(s) that is present in bacterial cells.

In *Staphylococcus aureus*, CDA synthesis and degradation are mediated by DacA (diadenylate cyclase) and GdpP (the primary CDA phosphodiesterase) respectively (1). Recent studies have highlighted that increased CDA concentrations promote tolerance to β-lactam antibiotics and allow cell wall restructuring (8, 11). Furthermore, a growing number of contemporary clinical surveillance studies have identified mutations in *gdpP* among β-lactam resistant or non-susceptible natural isolates of *S. aureus* (12–15). Since many of these mutations have been either shown (8) or are predicted to attenuate the function of GdpP, causing increased CDA concentrations in cells, these findings underscored the clinical importance of CDA and highlighted the importance of accurate determination of its concentrations for both basic and clinical research settings.

Detection and quantification of CDA are typically carried out either through HPLC/MS, indirect ELISA, or dye intercalation assay (16–18). These assays determine CDA’s abundance in a static manner, i.e. in samples containing bacterial cell lysates. Of these, HPLC/MS despite being a gold standard in the quantification of small molecules such as CDA requires expensive technical infrastructure and operational expertise. The operation of ELISA is relatively easy but requires expertise in protein purification. In this study, we present a novel RNA-based CDA biosensor (*bsuO* P6-4), which can determine its concentration in live cells through flow cytometric analysis.

*bsuO* P6-4 is a second-generation CDA biosensor with improved CDA affinity and signal-to-noise ratio than its predecessor, *yuaA* P1-4 (19). This improvement was achieved by fusing the pro-fluorescent dye-binding RNA aptamer (Spinach) to the P6 stem instead of the P1 stem (as in *yuaA* P1-4) of the natural CDA binding riboswitch sequence present in the upstream region of *ydaO/yuaA* genes in *Bacillus subtilis* (20) (Fig.1A). This rational design of *bsuO* P6-4 restored the pseudoknot interaction between the P1 and P8 stems, which acts as a native stabilizer of the *ydaO/yuaA* riboswitch structure (21–23). Additionally, a modified fluorescent dye binding module, coined as cpSpinach2 (24) that was circularly permutized to accept a transducer stem was used for the construction of *bsuO* P6-4 (Fig.1A). Thus, *bsuO* P6-4 consists of two components, the CDA-binding *ydaO/yuaA* module, and the pro-fluorescent, DFHBI-binding cpSpinach2 module. Binding of CDA to *bsuO* P6-4 enabled appropriate folding of the cpSpinach2 module, allowing DFHBI binding and production of a fluorescence signal (Fig.1B) and thereby enabling detection of CDA.

**Figure 1:**
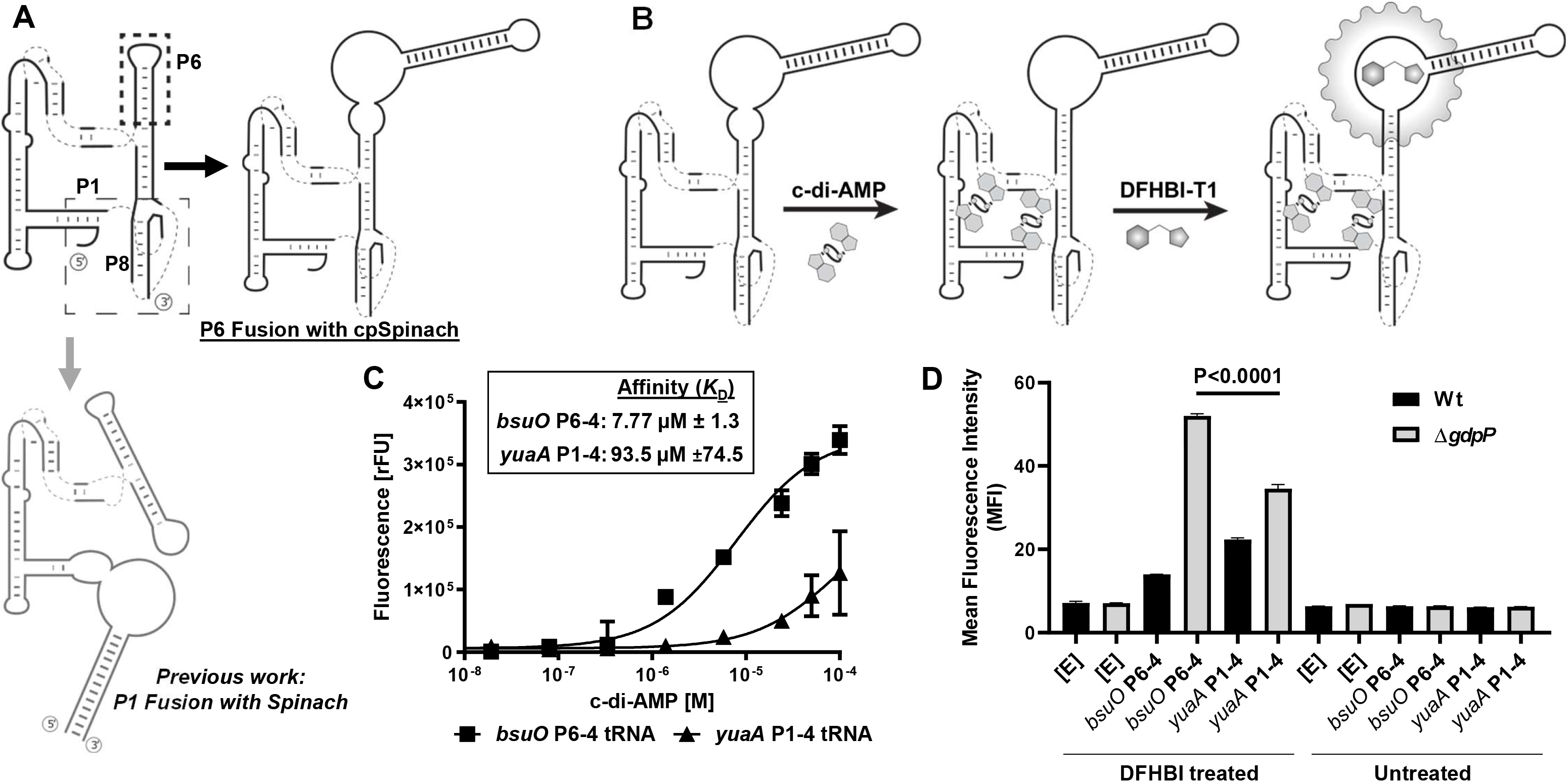
Construction of the cyclic-di-AMP biosensor *bsuO* P6-4 and its ability to detect cyclic-di-AMP. A) & B) Schematic representation of the creation and function of *bsuO* P6-4. C) & D) Comparison of *in-vitro* and *in-vivo* detection of CDA respectively by *bsuO* P6-4 and *yuaA* P1-4.

An *in-vitro* fluorescence assay testing the CDA biosensors revealed a >10-fold higher affinity of *bsuO* P6-4 compared to its predecessor *yuaA* P1-4 (Fig.1C). In preparation for *in-vivo* experiments, a tRNA scaffold was added to flank the 5’ and 3’ ends of the biosensors for increased RNA half-life (Table S1) (25). The resultant biosensors were cloned into a constitutive expression vector and transformed into a wild-type (Wt) *S. aureus* and its isogenic Δ*gdpP* strain. While both the biosensors were able to report the higher level of CDA that is characteristic of a Δ*gdpP* strain (8), *bsuO* P6-4 showed significantly enhanced fluorescence compared to that of *yuaA* P1-4. More importantly, *bsuO* P6-4 compared to *yuaA* P1-4 displayed a higher dynamic range of signal (3.73X vs 1.55X) between Wt and Δ*gdpP* strains making it amenable for detection of a wider concentration range of CDA (Fig.1D). The reason why *yuaA* P1-4 exhibits higher background fluorescence than *bsuO* P6-4 *in vivo* is unknown.

Next, we sought to determine whether *bsuO* P6-4 could report different concentrations of CDA in the cells. For this purpose, isogenic *gdpP* point mutants were created that displayed varying degrees of GdpP’s loss of function that were identified in our previous study (8). As shown in figure 2, *bsuO* P6-4 was able to detect differing CDA concentrations in the isogenic strains (Fig.2A), which correlated well (R^2^=0.9453) with the results independently obtained through ELISA assay among the identical strains (Fig.2B & C). In addition to the isogenic strains, *bsuO* P6-4 was also able to determine different CDA concentrations in clinical isolates (Fig. S1), which suggested that it could be used in both laboratory and clinical strains. Our results further showed that the Wt and Δ*gdpP* strains with *bsuO* P6-4 were also suitable for fluorescent microscopic analysis (Fig. S2). However, this method was not sensitive enough to differentiate the intermediate concentrations of CDA of the other isogenic strains (data not shown).

**Figure 2:**
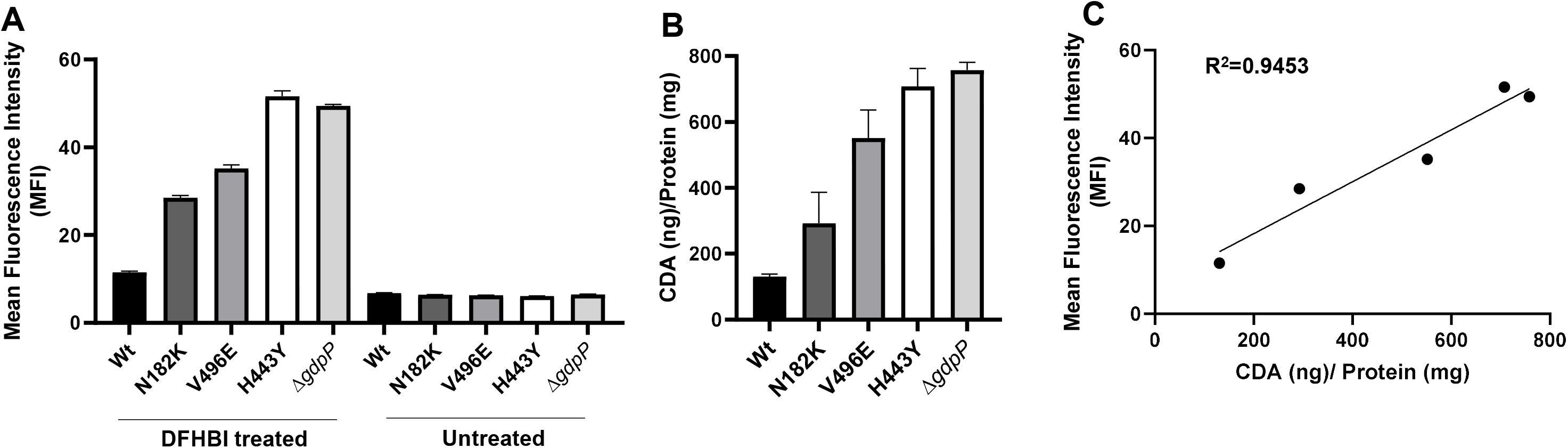
Ability of *bsuO* P6-4 in detecting varying concentrations of cyclic-di-AMP in *S. aureus*. A) & B) Detection of CDA in wild-type and isogenic strains of *S. aureus* which carried GdpP loss of function mutations using flow cytometry and ELISA assay respectively. C) Correlation of signals obtained in A & B.

In summary, we have developed a novel *bsuO* P6-4 biosensor that is effective in determining different CDA concentrations in live *S. aureus* cells through flow cytometry. The plasmid harboring *bsuO* P6-4 can be transformed into both laboratory and clinical *S. aureus* isolates for reporting CDA concentration. This biosensor-based approach could be used in flux detection of CDA concentrations in future studies.

## ACKNOWLEDGMENTS

This work was supported by NIH grants 2R01AI100291 and startup funds provided by the University of Maryland, Baltimore, and the University of Maryland Center for Environmental Science to SSC and NIH grant R01 GM124589 to MCH. The authors would also like to thank the Charles A. and Lois H. Miller Foundation for their generous gift to purchase the flow cytometer used in this study.

