## Supplementary material for "*In-vivo* detection of cyclic-di-AMP in *Staphylococcus aureus*"

**Running Title**: RNA biosensor based detection of cyclic-di-AMP in *S. aureus*

Nagaraja Mukkayyan,[a,b,#] Raymond Poon,[a,b,#] Philipp N. Sander,[c] Li-Yin Lai,[a,b] Zahra Zubair-Nizami,[a,b] Ming C. Hammond[c,d]* and Som S. Chatterjee[a,b,e]*

[a] Department of Microbial Pathogenesis, School of Dentistry, University of Maryland, Baltimore, MD.

[b] Institute of Marine and Environmental Technology, Baltimore, MD.

[c] Department of Chemistry, University of California, Berkeley, CA.

[d] Department of Chemistry and Henry Eyring Center for Cell and Genome Sciences, University of Utah, Salt Lake City, UT.

[e] University of Maryland Center for Environmental Science, Baltimore, MD.

[#] Equal contributors

### MATERIALS AND METHODS:

#### Bacterial strains, growth media, and growth conditions

Bacterial strains used in this study are shown in Table S2. Bacteria were grown in trypticase soy broth (TSB) at 37°C with shaking at 180 rpm or on trypticase soy agar (TSA) plates at 37°C. Strains containing plasmids *pTX*Δ, *pJB*38 and *pET*28b(+) were grown in tetracycline (12.5 µg/mL), chloramphenicol (10 µg/mL), and kanamycin (50 µg/mL) respectively as a selection markers. The molecular cloning experiments were carried out using *S. aureus* RN4220 strains. Lists of plasmids and primers used in this study are shown in Table S3 and S4 respectively. Sequence fidelity of all the mutants and constructs was validated using analytical PCR and Sanger sequencing.

#### Construction of mutants

Construction of mutants with *gdpP* point mutations (i.e. N182K, V496E, and H443Y) was carried out using plasmid *pJB*38 as described previously (1, 2). Briefly, a splice-overlap PCRs of the 1 kb up- and down-stream genomic region surrounding the *gdpP* mutations were amplified using either primers gdpP-SacI1-Nterm-F and gdpP-XmaI-Nterm-R or gdpP-SacI-Cterm-F and gdpP-XmaI-Cterm-R and with the primers labeled with the respective point mutations (Table S4). The resulting PCR product was digested with *SacI* and *XmaI* and ligated into a *pJB38*. The plasmid was transformed into SF8300 and the standard allelic replacement procedure was carried out as shown before (1). The mutants were sequence verified to confirm the *gdpP* point mutations.

#### Construction of *bsuO* P6-4, *In-vitro* transcription, *in-vitro* CDA affinity determination, and cloning into *pTX*Δ

a) Construction of *bsuO* P6-4: *bsuO* P6-4 was constructed by Golden Gate cloning. Briefly, the *Bacillus subtilis ydaO* (*bsuO*) sequence cloned into a *TOPO PCR 2.1* vector was PCR-amplified “around-the-horn” with overhang primers BsuO-P6-Fwd and BsuO-P6-Rev to add *BsaI* restriction sites at the P6 stem sequence. A circularly permutized Spinach2 sequence (cpSpinach2 (3) was amplified with overhang primers BsuO-P6-4-Spin-Fwd and BsuO-P6-4-Spin-Rev adding sticky ends and *BsaI* restriction sites. After golden gate assembly (restriction with *BsaI*-HF, 50°C, 1h, and ligation with T4 ligase, 30 min, 24°C, both NEB), constructs were transformed into TOP10 chemically competent *E. coli* (Invitrogen) and clones recovered and sequenced. To assemble tRNA construct via Gibson Assembly, *pET-31b tRNA* (4) was amplified with primers adding homology sites to *bsuO* (BsuO-tRNA-GA-Fwd and BsuO-tRNA-GA-Rev), and *bsuO* P6-4 was amplified with primers adding homology sites to the tRNA scaffold (tRNA-BsuO-GA-Fwd and tRNA-BsuO-GA-Rev). After Gibson assembly (NEB), constructs were transformed into TOP10 chemically competent *E. coli* (Invitrogen) and clones recovered and sequenced.

b) *In-vitro* transcription: Biosensor RNA was transcribed using T7 RNA polymerase (NEB) and previously published protocols (5). In brief, a DNA transcription template was produced by PCR amplification with primers T7-Fwd and tRNA-Rev from a sequence-confirmed plasmid PCR template. Transcribed products were purified by PAGE, quantified after thermal hydrolysis (6), and stored at -80°C.

c) CDA affinity determination: Apparent affinity ($K_d$) was determined by fluorescence response in a 384-well plate reader assay, as published previously. Biosensor RNA (30 nM), DFHBI (10 µM), were incubated reaction buffer (40 mM HEPES, pH 7.5, 125 mM KCl, 3 mM $MgCl_2$) with increasing concentrations of CDA (Biolog) at 37°C. Upon reaching equilibrium after 3h, fluorescence (Excitation 448/Emission 508 nm) was determined using a SpectraMax Paradigm plate reader and analyzed using GraphPad Prism 9.

d) Cloning into *pTX*$_{\Delta}$: Biosensor constructs were PCR amplified using primers Spinach-BamHI-F and Spinach-MluI-R (Table S4), digested with *BamHI* and *MluI*, and were ligated into the *pTX*$_{\Delta}$ plasmid. The resulting plasmid was transformed into RN4220 via electroporation and were sequence verified. The plasmid was transduced into the final strains using the phage Φ11.

**Quantification of c-di-AMP levels in *S. aureus using* flow cytometer.**

Quantification of CDA was carried out as previously described in Kellenberger et al., 2015 (7), with slight modifications. *S. aureus* strains containing *pTX*$_{\Delta}$+*busO* P6-4, *pTX*$_{\Delta}$+*yuaA* P1-4, or an empty *pTX*$_{\Delta}$ vector were inoculated using a 1 µL loop from a plate into culture tubes containing 4 mL of TSB media containing tetracycline (12.5 µg/mL) and shaken at 37°C for 18-22 h at 180 rpm. 500 $OD_{600}$ worth of bacterial cultures were harvested and pelleted by centrifugation at 6000 xg for 5 min. The cell pellets were washed with 1x PBS and then resuspended in 100 µl of TSB media containing tetracycline (12.5 µg/mL) and with or without DFHBI-1T (200 µM). The resuspended bacterial samples were incubated at 37ºC for 1 h in the dark. After incubation, 40 µL of bacteria was diluted into 1 mL of 1x PBS, 200 µl of which was dispensed into a 96-well plate in triplicate, and analyzed with a Guava easyCyte Flow cytometer (Luminex, USA) (Parameters: 30,000 events, Excitation: Blue laser (488 nm), Emission: Green channel (512-530 nm), Fluidics: Slow, Cut-off: 0). The data was analyzed using InCyte from guavaSoft 4.0 software and the mean fluorescent intensity from the GFP channel was calculated for each sample.

**Quantification of cyclic-di-AMP levels in *S. aureus* using competitive ELISA assay**

Determination of CDA levels using Competitive ELISA assay was carried out as described in the Underwood *et. al.,* 2014 (8), with slight modifications.

a) CabP cloning and Protein purification: The *cabP* gene was PCR amplified from *Streptococcus pneumoniae* D39 genomic DNA using primers cabP-NdeI-F and cabP-HindIII-R and cloned into the *pET28b(+)* vector using *NdeI* and *HindIII* restriction enzymes. The resulting plasmid was transformed into DH5α via heat shock and sequence-verified for accuracy. The resulting *pET28b(+)* + *cabP* was transformed into *E. coli* BL21(DE3). CabP protein was overexpressed and purified as described in Bai *et. al.,* 2014 (9).

b) CDA extraction from *S. aureus*: Bacteria were inoculated using a 1 uL loop from a plate into culture tubes containing 4 mL of TSB media and shaken at 37°C for 18-22h at 180 rpm. 30 $OD_{600}$ of bacterial culture was harvested, pelleted at 2739 xg for 10 min, and washed with 1x PBS. Cell pellets were resuspended with 600ul of 50 mM Tris HCl (pH 8.0) and lysed using a FastPrep-24 set to 6.5 m/s and 45 seconds for 4 cycles (MP Biomedicals). Lysed cells were centrifuged at 17000xg for 10mins at 4°C and supernatants were collected. An aliquot of the total lysate was saved and later used for protein estimation using Pierce BCA Protein Assay kit (Thermo Fisher). Each sample's CDA concentration (ng/mL) was normalized with respect to protein content (mg/mL). Supernatants were boiled at 95°C for 10 mins, allowed to rest to room temperature for 10 min, and centrifuged at 17000 xg for 10mins. The boiled supernatants were collected, diluted in half with 50 mM Tris HCl (pH 8.0), and used for Competitive ELISA assay for CDA quantification.

c) Competitive ELISA assay: CDA quantification carried out as described in Underwood *et. al.,* 2014 (8). HRP-conjugated streptavidin (Thermo Scientific) diluted to a 1:5000 dilution in PBS was used after incubation of samples on the ELISA plate.

#### Fluorescence Microscopy

*S. aureus* strains were treated similarly as described above for flow cytometry assay. 3µl of the DFHBI-1T incubated bacterial samples were smeared onto a glass slide and air-dried for 10min at room temperature in the dark. Microscopic analysis was carried out with a Zeiss Axio Observer Z1 Inverted microscope (Carl Zeiss. Inc. Germany). DIC and GFP images were captured using a 100X oil immersion objective. The images were analyzed using Volocity software version 6.5.1 (Quorum Technologies).

#### Statistical Analysis

Statistical analysis was carried out using GraphPad Prism 8.1.1. Comparisons and significance were determined using Student's t-test. Each experiment was repeated at least twice to ensure reproducibility.

### SUPPLEMENTARY TABLES AND FIGURES:

**Table S1:** Biosensor sequences used in this study.

| Construct | Spinach variant | Sequence: 5' -> 3' |
|---|---|---|
| *bsuO* P6-4 | cpSpinach | GGaaucgcuuaaucugaaaucagagcgggggacccaauagaacggcuuuuugccguuggggugaauccu**uuguugaguagagugugagcuccguaacuaguuacaucgcaagauguaacugaaugaaauggugaaggacgguucca**agguagggcuaacucucauaugcccgaauccgucagcuaaccucguaagcguucgugagaggaUC |
| *yuaA* P1-4 | Spinach2 | **GAUGUAACUGAAUGAAAUGGUGAAGGACGGGUCCA**gcuuaaucaaacacgaacgggggaaccaacgauuggcuguuuuauuaacagccuuggggugaaucuuacuaaguaagagggguacucugaaucccuaauccgacagcuaaccucguaggc**UUGUUGAGUAGAGUGUGAGCUCCGUAACUAGUUACAUC** |
| *bsuO* P6-4 tRNA | cpSpinach | *GCCCGGAUAGCUCAGUCGGUAGAGCAGCGGCCGGAAAACA*aaucgcuuaaucugaaaucagagcgggggacccaauagaacggcuuuuugccguuggggugaauccu**uuguugaguagagugugagcuccguaacuaguuacaucgcaagauguaacugaaugaaauggugaaggacgguucca**agguagggcuaacucucauaugcccgaauccgucagcuaaccucguaagcguucgugagaggaGAUGCGGCCGCGGGUCCAGGGUUCAAGUCCCUGUUCGGGCGCCA |
| *yuaA* P1-4 tRNA | Spinach2 | GCCCGGAUAGCUCAGUCGGUAGAGCAGCGGCCGG**AUGUAACUGAAUGAAAUGGUGAAGGACGGGUCCA**gcuuaaucaaacacgaacgggggaaccaacgauuggcuguuuuauuaacagccuuggggugaaucuuacuaaguaagagggguacucugaaucccuaauccgacagcuaaccucguaggc**UUGUUGAGUAGAGUGUGAGCUCCGUAACUAGUUACAUC**CGGCCGCGGGUCCAGGGUUCAAGUCCCUGUUCGGGCGCCA |

**Bold** sequences indicate the Spinach2 or cpSpinach2 sequence. The tRNA scaffold is underlined.

**Table S2**: List of strains used in this study.

| Strain | Description | Reference |
|---|---|---|
| RN4220 | Laboratory *S. aureus* strain | 2 |
| SF8300 Wt | *S. aureus* USA300 MRSA clinical isolate | " |
| SF8300 Δ*gdpP* | *gdpP* deleted SF8300 strain | " |
| SF8300 [E] | SF8300 with empty constitutively expression vector $pTX_{\Delta}$ | " |
| SF8300 Δ*gdpP* [E] | SF8300 Δ*gdpP* with empty constitutively expression vector $pTX_{\Delta}$ | " |
| Strain 1 | *S. aureus* ST3405/CC8 lineage strain with *gdpP* mutation V496E | " |
| Strain 9 | *S. aureus* ST3412/CC101 lineage strain with *gdpP* mutation E486K | " |
| SF8300 [*bsuO* P6-4] | SF8300 with $pTX_{\Delta}$ expressing RNA-aptamer *bsuO* P6-4 | This study |
| SF8300 [*yuaA* P1-4] | SF8300 with $pTX_{\Delta}$ expressing RNA-aptamer *yuaA* P1-4 | " |
| SF8300 Δ*gdpP* [*bsuO* P6-4] | SF8300 Δ*gdpP* with $pTX_{\Delta}$ expressing RNA-aptamer *bsuO* P6-4 | " |
| SF8300 Δ*gdpP* [*yuaA* P1-4] | SF8300 Δ*gdpP* with $pTX_{\Delta}$ expressing RNA-aptamer *yuaA* P1-4 | ` |
| SF8300 *gdpP* (N182K) | SF8300 with *gdpP* mutation N182K | " |
| SF8300 *gdpP* (V496E) | SF8300 with *gdpP* mutation V496E | " |
| SF8300 *gdpP* (H443Y) | SF8300 with *gdpP* mutation H443Y | " |
| SF8300 *gdpP* (N182K) [E] | SF8300 *gdpP* (N182K) with empty constitutively expression vector $pTX_{\Delta}$ | " |
| SF8300 *gdpP* (N182K) [*bsuO* P6-4] | SF8300 *gdpP* (N182K) with $pTX_{\Delta}$ expressing RNA-aptamer *bsuO* P6-4 | " |
| SF8300 *gdpP* (V496E) [E] | SF8300 *gdpP* (V496E) with empty constitutively expression vector $pTX_{\Delta}$ | " |
| SF8300 *gdpP* (V496E) [*bsuO* P6-4] | SF8300 *gdpP* (V496E) with $pTX_{\Delta}$ expressing RNA-aptamer *bsuO* P6-4 | " |
| SF8300 *gdpP* (H443Y) [E] | SF8300 *gdpP* (H443Y) with empty constitutively expression vector $pTX_{\Delta}$ | " |
| SF8300 *gdpP* (H443Y) [*bsuO* P6-4] | SF8300 *gdpP* (H443Y) with $pTX_{\Delta}$ expressing RNA-aptamer *bsuO* P6-4 | " |
| Strain 1 [E] | Strain 1 with empty constitutively expression vector $pTX_{\Delta}$ | " |
| Strain 1 [*bsuO* P6-4] | Strain 1 with $pTX_{\Delta}$ expressing RNA-aptamer *bsuO* P6-4 | " |
| Strain 9 [E] | Strain 9 with empty constitutively expression vector $pTX_{\Delta}$ | " |
| Strain 9 [*bsuO* P6-4] | Strain 9 with $pTX_{\Delta}$ expressing RNA-aptamer *bsuO* P6-4 | " |
| BL21(DE3) *pET28b(+)* + *cabP* | *E. coli* with *pET28b(+)* expression system for CabP Protein | " |

**Table S3**: List of plasmids used in this study.

| Plasmid | Description | Reference |
|---|---|---|
| *pTX*$_{\Delta}$ | Empty plasmid for constitutive expression | 2 |
| *pTX*$_{\Delta}$ + *yuaA* P1-4 | Constitutively expressed RNA-aptamer *yuaA* P1-4 | This Study |
| *pTX*$_{\Delta}$ + *bsuO* P6-4 | Constitutively expressed RNA-aptamer *bsuO* P6-4 | " |
| *pJB38* + *gdpP* (N182K) | Introduce N182K mutation into *gdpP* in SF8300 | " |
| *pJB38* + *gdpP* (V496E) | Introduce V496E mutation into *gdpP* in SF8300 | " |
| *pJB38* + *gdpP* (H443Y) | Introduce H443Y mutation into *gdpP* in SF8300 | " |
| *pET28b(+)* + *cabP* | CabP protein overexpression vector | " |

**Table S4**: List of primers used in this study.

| Primer | Sequence (5'- 3') | Description |
|---|---|---|
| Spinach-BamHI-F | AAAGGATCCGCCCGGATAGCTCAGTCGGTAGAGC | *bsuO* P6-4 and *yuaA* P1-6 Cloning in *pTX*Δ |
| Spinach-MluI-R | TTTACGCGTCAAAAAACCCCTCAAGACCCGTTTAGAGG | " |
| gdpP-SacI1-Nterm-F | AGTAGCGATGAGCTCTAATTTCATTAATCGCTTTTATG | SF8300 *gdpP*(N182K) mutation creation |
| gdpP-XmaI-Nterm-R | AAAACCAGTTCCCGGGTTTATGCGTATCAAC | " |
| gdpP-SacI-Cterm-F | TAGATGGTTTGAGCTCTCAAATTTCAACAAC | SF8300 *gdpP*(V496E) and *gdpP*(H443Y) mutation creation |
| gdpP-XmaI-Cterm-R | TCGTATATCCCGGGGAATGAATTCCATTTGG | " |
| gdpP-N182K-F | ATTATTTTTAGATAAATACGATGAGATTACGCAAAATATG | SF8300 *gdpP*(N182K) mutation creation |
| gdpP-N182K-R | TCTCATCGTATTTATCTAAAAATAATGTCGCAATGATTGG | " |
| gdpP-H443Y-F | TTGTTATCGATCATTATAGACGTGGTGAAAGCTTCATCTC | SF8300 *gdpP*(V496E) mutation creation |
| gdpP-H443Y-R | ACCACGTCTATAATGATCGATAACAACTTTACGGTTTGC | " |
| gdpP-V496E-F | ATGCAGGTATTATTGAAGATACAAGAAACTTTACATTACG | SF8300 *gdpP*(H443Y) mutation creation |
| gdpP-V496E-R | AAGTTTCTTGTATCTTCAATAATACCTGCATACATCACTG | " |
| cabP-NdeI-F | TTTCATATGTCAGATCGTACGATTGG | *cabP* Cloning in pET28b(+) |
| cabP-HindIII-R | TTTAAGCTTACGAATTCAATGCTAC | " |
| BsuO-P6-Fwd | TTAGGTCTCTAGGGCTAACTCTCATATGC | Attach 3' *BsaI* cut site at the P6 stem root of *bsuO* |
| BsuO-P6-Rev | TTAGGTCTCATTCACCCCAACGG | Attach 5' *BsaI* cut site at the P6 stem root of *bsuO* |
| BsuO-P6-4-Spin-Fwd | ATGGTCTCGTGAATCCTTTGTTGAGTAGAGTGTG | Attach 5' *BsaI* cut site to cpSpinach |
| BsuO-P6-4-Spin-Rev | TGGTCTCGCCCTACCTTGGACCCGTCC | Attach 3' *BsaI* cut site to cpSpinach |
| tRNA-BsuO-GA-Fwd | AGTCGGTAGAGCAGCGGCCGGAAAACAAATCGCTTAATCTGAAATC | Gibson Assembly primer adding 5' tRNA scaffold homology to *bsuO* sequence |
| tRNA-BsuO-GA-Rev | GAACCCTGGACCCGCGGCCGCATCTCCTCTCACGAACGCT | Gibson Assembly primer adding 3' tRNA scaffold homology to *bsuO* sequence |
| BsuO-tRNA-GA-Fwd | AGCGTTCGTGAGAGGAGATGCGGCCGCGGGTCCAGGGTTC | Gibson Assembly primer adding 5' *bsuO* scaffold homology to tRNA sequence |
| BsuO-tRNA-GA-Rev | AGATTAAGCGATTTGTTTTCCGGCCGCTGCTCTACCGACT | Gibson Assembly primer adding 3' *bsuO* scaffold homology to tRNA sequence |
| T7-Fwd | CGAAATTAATACGACTCACTATAGGG | Transcription Template 5' primer |
| tRNA-Rev | TGGCGCCCGAACA | Transcription Template 3' primer |

**Figure S1:** A) & B) Detection of CDA in wild-type and clinical strains of *S. aureus* which carried GdpP loss of function mutations using flow cytometry and ELISA assay respectively. C) Correlation of signals obtained in A & B.

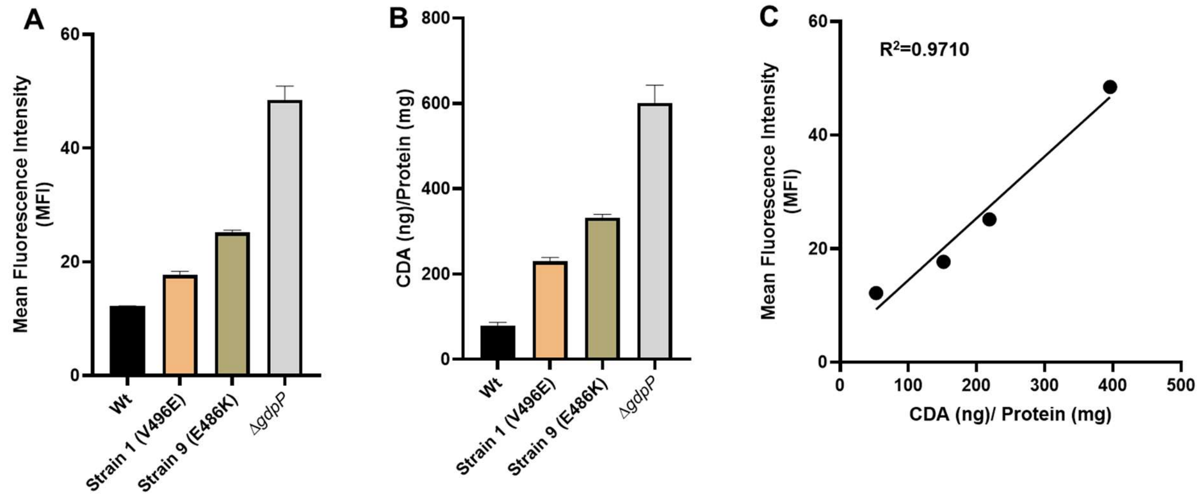

**Figure S2:** Bright field (Upper panels) and fluorescent (lower panels) microscopic images of *S. aureus* wild-type and Δ*gdpP* strain expressing *bsuO* P6-4. Scale bar in the images represent 5μm.

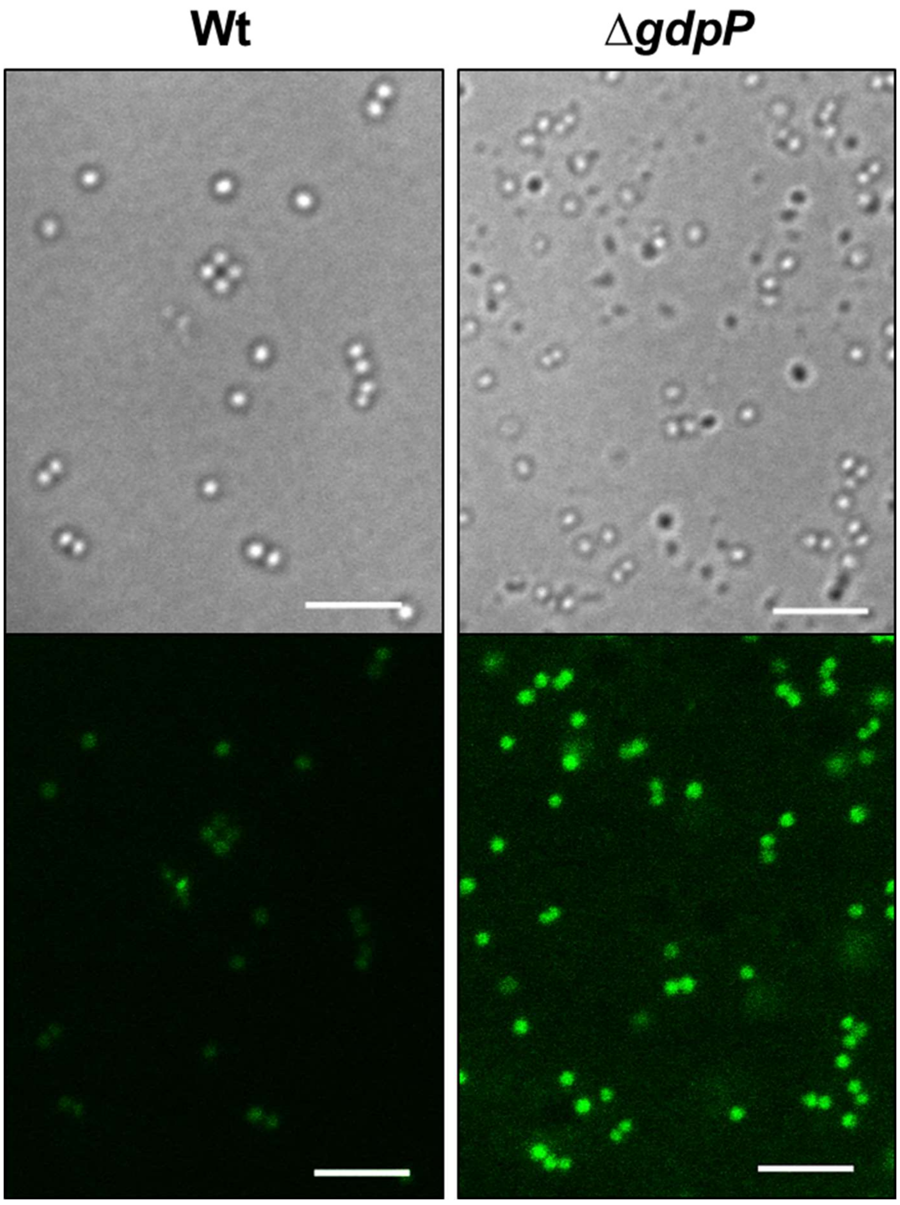
